# The unsolved problem of otitis media in indigenous populations: A systematic review of upper respiratory and middle ear microbiota in indigenous children

**DOI:** 10.1101/355982

**Authors:** Andrea Coleman, Amanda Wood, Seweryn Bialasiewicz, Robert S Ware, Robyn L Marsh, Anders Cervin

## Abstract

**Background:** Otitis media (OM) imposes a great burden of disease in indigenous populations around the world, despite a variety of treatment and prevention programs. Improved understanding of the pathogenesis of OM in indigenous populations is required to advance treatment and reduce prevalence. We conducted a systematic review of the literature exploring upper airway and middle ear microbiota in relation to OM in indigenous children.

**Methods:** Papers targeting microbiota in relation to OM in children <18 years indigenous to Australia, New Zealand, North America, and Greenland were sought. MEDLINE, CINAHL, EMBASE, Cochrane Library, and Informit databases were searched using key words. Two independent reviewers screened titles, abstracts, and then full-text papers against inclusion criteria according to PRISMA guidelines.

**Results:** Twenty-five papers considering indigenous Australian, Alaskan and Greenlandic children were included. There were high rates of nasopharyngeal colonization with the three main otopathogens (*Haemophilus influenzae*, *Streptococcus pneumoniae*, and *Moraxella catarrhalis*) in indigenous children with OM. Middle ear samples had lower rates of otopathogen detection, although detection rates increased when molecular methods were used. *Pseudomonas aeruginosa* and *Staphylococcus aureus* were commonly detected in middle ear discharge of children with chronic suppurative OM. There was significant heterogeneity between studies, particularly in microbiological methods, which were largely limited to culture-based detection of the main otopathogens.

**Conclusions:** There are high rates of otopathogen colonization in indigenous children with OM. Chronic suppurative OM appears to be associated with a different microbial profile. Beyond the main otopathogens, the data are limited. Further research is required to explore the entire upper respiratory tract/ middle ear microbiota in relation to OM, with the inclusion of healthy indigenous peers as controls.

## Introduction

Otitis media (OM) describes a spectrum of pathologies that involve inflammation and/or infection in the middle ear. This spectrum encompasses a continuum from acute to chronic disease that is clinically characterised by fluid in the middle ear [1–4]. OM is highly prevalent in indigenous populations globally, particularly when compared to non-indigenous peers [5, 6], and often occurs earlier, more frequently and in more severe forms [4, 5, 7]. Prevalence data reports that up to one-third of Greenlandic and Alaskan Inuit, Native American and Australian Indigenous children suffer from chronic suppurative OM (CSOM) [6, 8–11]. The World Health Organization considers CSOM prevalence of ≥4% indicative of a public health problem serious enough to require urgent attention [12]. OM-related complications result in approximately 21,000 deaths each year worldwide [13]. OM-associated hearing loss can impact significantly on language and social skills development, school attendance and educational outcomes, and downstream effects such as greater contact with the criminal justice system later in life [4, 14, 15]. Medical interventions including liberal antibiotic prescription and vaccination programs have limited effectiveness in indigenous populations [16–19], thus new treatment avenues need to be considered.

The reasons for high OM prevalence in indigenous populations are likely to be multi-factorial. Risk factors include poverty, inadequate housing, overcrowding, and exposure to environmental tobacco smoke [6, 8, 20, 21]. These risk factors are ubiquitous across indigenous populations worldwide [22]. Genetic susceptibility to OM has not been studied in indigenous populations [23, 24].

The microbiota of the upper respiratory tract (URT) is an important OM risk factor across all populations. Most research to date has focused on the role of the three main otopathogens: *Streptococcus pneumoniae*, *Moraxella catarrhalis*, and non-typeable *Haemophilus influenzae* [25]. It is not currently clear whether commensal bacteria amongst the URT microbiota contribute to, or mitigate, OM risk in indigenous children. In non-indigenous children, 16S ribosomal RNA (rRNA) gene analyses have suggested that a ‘healthy’ nasopharyngeal (NP) microbiota is more diverse than that of children with OM [26–29]. This ‘healthy’ NP microbiota contains bacteria that may be protective or promote microbiota stabilization, including, *Moraxella, Corynebacterium, Dolosigranulum, Propionibacterium (Cutibacterium)*, *Lactococcus and Staphylococcus* [26–29]. It is currently unknown whether these results are generalizable to indigenous populations.

While high rates of OM are reported for many developing countries, indigenous populations, as defined by the United Nations [30], share unique challenges in relation to OM. The aim of this systematic review is to assess the data pertaining to the NP and middle ear microbiota in relation to OM affecting indigenous children around the world.

## Methods

Methods used for this systematic review were developed with reference to the Preferred Reporting Items for Systematic Reviews and Meta-Analyses (PRISMA) Statement. The protocol was registered with the International Prospective Register of Systematic Reviews (PROSPERO) (CRD42016033905) prior to commencement.

### Inclusion criteria

All studies exploring the microbiota of the URT (nose, nasopharynx, mouth, oropharynx, throat, tonsils, adenoid, and middle ear) in relation to OM in indigenous children aged 0-18 years old were included. For studies that included children without OM and/or did not report microbiology results specifically for children with OM, either only middle ear data were included, or if only the NP was sampled, the studies were excluded. Indigenous populations from Australia, New Zealand, Unites States of America, Canada, and Greenland were included.

### Search strategy

Literature search strategies were developed in collaboration with a health sciences librarian using medical subject headings (MeSH) and key words (Additional File 1). The following electronic databases were searched from inception until 15 August 2017: MEDLINE (from 1946) and CINHAL (from 1982) via EBSCOhost, EMBASE (from 1966), Cochrane Library (from 1996), and Informit (from 1990-April). To ensure search saturation, we reviewed the reference lists of relevant studies and sought unpublished clinical audits through the Australian Institute of Health and Welfare (https://www.aihw.gov.au/) and The Australian Indigenous Health Info Net (https://healthinfonet.ecu.edu.au/). Two independent reviewers (ACol and AW) revised titles and abstracts, then full text publications with reference to the inclusion criteria. Study selection interrater agreement between the two reviewers was calculated as the proportion of positive agreement (PA) [31].

### Data extraction

Two independent reviewers (ACol and AW) extracted data in duplicate onto a Microsoft Excel spreadsheet. Publication authors were contacted where data had been represented graphically or data were missing. We screened for multiple reports from the same study, and where multiple reports existed, compared and extracted relevant data; if inconsistencies existed, we contacted the authors for clarification. The following data were extracted for all studies meeting inclusion criteria: publication year, geographical location, study design, number of participants, age range, ethnicity, number of participants with an OM diagnosis, type of OM, number of controls, anatomical location of sample(s), microbiota investigation method, type and quantity of bacteria, viruses, and fungi detected from each anatomic site. For the purpose of the review ‘culture’ is defined as culture targeting the three main otopathogens and ‘extended culture’ is defined as culture used to detect bacteria beyond these otopathogens. Only quantitative PCR (qPCR) data were included when both culture and qPCR were used. For longitudinal studies, data relating to both the number of swabs and number of children were extracted, when there were multiple swabs per child. For data obtained from clinical trials, we included data only from samples collected prior to randomization.

### Data analysis

Where there were a sufficient number of studies, meta-analysis of proportions were calculated using random effects analysis via Stata/IC 15, otherwise we synthesized the data into a systematic narrative. We calculated heterogeneity using I^2^ statistic.

### Risk of bias assessment

Two independent reviewers (ACol and AW) assessed the risk of bias for each study with reference to the Critical Appraisal Skills Program (CASP) Cohort Study Checklist [32]. Within the CASP Checklist, we assessed for the following confounding variables: age, overcrowding, antibiotic use, daycare/ school attendance, and concurrent respiratory/ upper respiratory tract infection. Study quality was categorized as ’poor’, ’moderate’ or ’good’ based on the CASP Checklist. The overall quality of evidence was judged as high, moderate, low, and very low [33].

## Results

The initial search identified 5592 articles. After screening titles, the abstracts of 956 articles and 332 full text publications were reviewed (Figure 1). There was substantial PA between the reviewers of titles (PA = 0.68) and abstracts (PA = 0.79). Twenty-five articles met the inclusion criteria; these were from Australian Indigenous (n=22), Greenlandic (n=2) and Alaskan Inuit (n=1). No papers reported OM otopathogens or microbiota in Native American or New Zealand Maori children.

**Figure 1:**
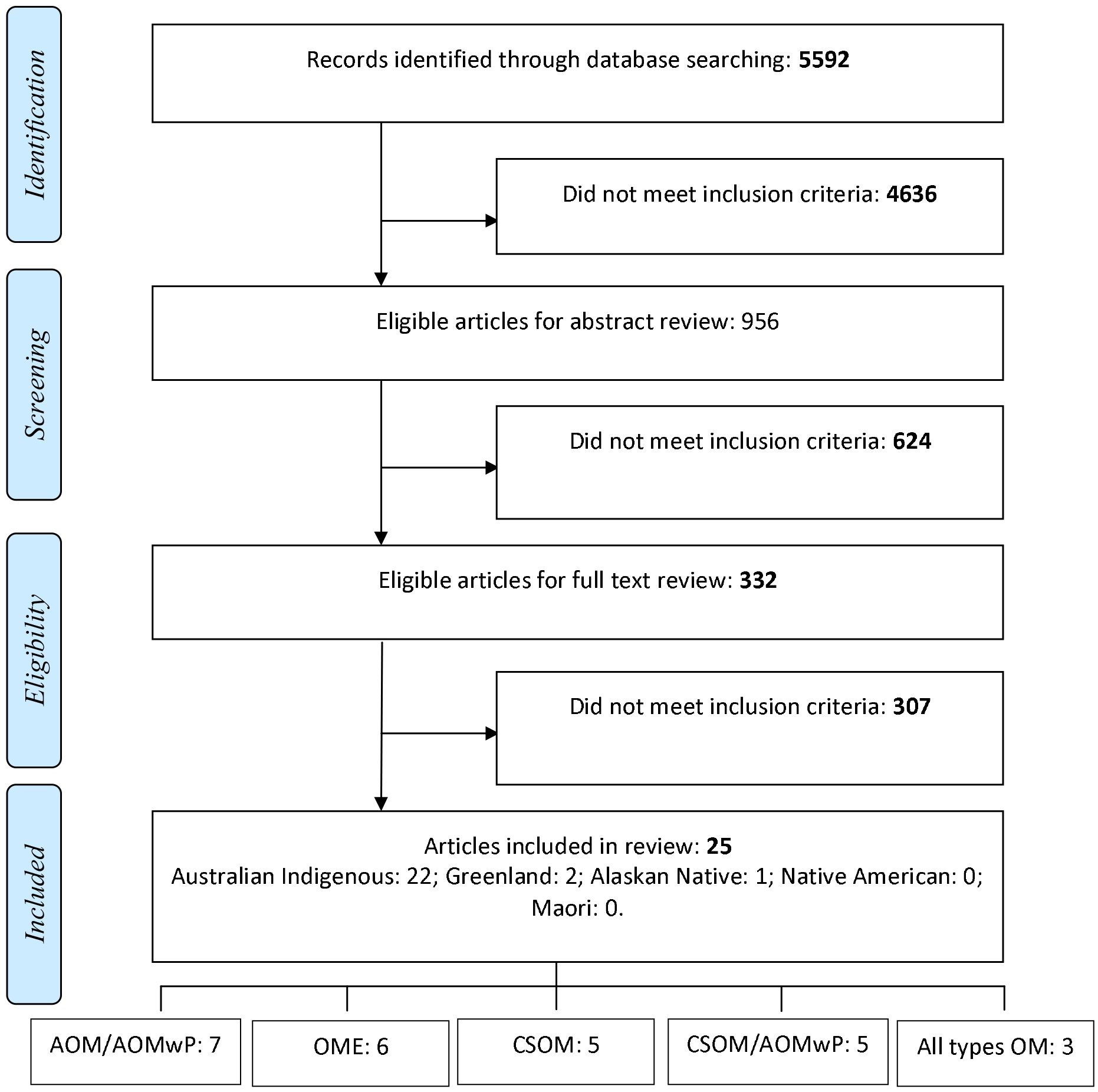
Literature search and selection.

### Risk of bias assessment

According to the CASP risk of bias assessment, most studies (80%) were judged as either ‘poor’ or ‘moderate’, largely due to confounding variables not being considered (Table 1). Recruitment bias was difficult to assess, as recruitment processes were often poorly documented. Only one study [34] included healthy indigenous controls and another three [21, 35, 36] included children without OM enrolled in longitudinal studies. Within indigenous populations, participants were recruited from limited geographical regions, making generalization beyond these regions difficult. Overall, the quality of the literature was ‘low’.

**Table 1:** Risk of bias assessment.

|  | Reference | Focused aim? | Cohort recruitment acceptable? | Exposure accurately measured? | Outcome accurately measured? | Confounding factors identified? | Confounding factors accounted for? | Are the results precise? | Are the results believable? | Do results fit with other available data | Overall quality score |
| --- | --- | --- | --- | --- | --- | --- | --- | --- | --- | --- | --- |
| 1972, Stuart* | [46] | + | + | - | - | + | - | - | - | - | Poor |
| 1975, Copeman | [44] | - | - | - | - | - | - | ? | - | - | Poor |
| 1975, Stuart | [45] | + | + | - | - | - | - | - | - | - | Poor |
| 1985, Dawson | [55] | + | - | + | + | - | - | ? | - | - | Poor |
| 1994, Leach | [21] | + | + | + | + | - | - | - | + | + | Mod |
| 1996, Homøe | [34] | + | + | + | + | + | + | + | + | + | Good |
| 1999, Parkinson | [38] | + | + | + | + | - | - | + | + | + | Mod |
| 2003, Couzos* | [40] | + | ? | + | + | + | + | + | + | + | Good |
| 2003, Stuart | [49] | + | ? | + | + | + | - | + | + | + | Mod |
| 2005, Gibney | [50] | + | + | + | ? | - | - | + | + | + | Poor |
| 2006, Leach* | [51] | + | ? | + | + | - | - | - | - | + | Poor |
| 2007, Ashhurst-Smith | [37] | + | ? | + | + | - | - | ? | + | - | Mod |
| 2008, Leach* | [43] | + | + | + | + | - | - | + | + | + | Mod |
| 2008, Leach* | [54] | + | + | + | + | - | - | + | + | + | Poor |
| 2009, Homøe* | [39] | + | + | + | + | - | - | - | - | - | Poor |
| 2009, Mackenzie* | [52] | + | ? | - | + | - | - | + | - | + | Poor |
| 2010, Morris * | [53] | + | + | + | + | + | + | + | + | + | Good |
| 2011, Binks | [35] | + | ? | ? | + | - | - | + | + | + | Poor |
| 2012, Marsh | [56] | + | ? | + | + | - | - | + | + | + | Mod |
| 2012, Sun | [36] | + | + | + | + | + | + | + | + | + | Good |
| 2013, Smith-Vaughan | [48] | + | + | + | + | - | - | + | + | + | Mod |
| 2013, Stephen* | [42] | + | ? | + | + | - | - | + | + | + | Mod |
| 2015, Jervis-Bardy | [57] | + | ? | + | + | + | + | + | + | + | Good |
| 2015, Leach* | [47] | + | ? | + | + | - | - | + | + | + | Mod |
| 2016, Leach* | [17] | + | ? | + | + | + | - | + | + | + | Mod |

Data based on CASP-based risk of bias assessment. Assessment of bias pertained to the
microbiology data, and not to clinical data.
\* Indicates studies where microbiological outcomes were not the primary outcome. ? indicates
that this variable was unable to be assessed.

### Heterogeneity

The literature was limited by methodological and statistical heterogeneity across the studies, including heterogeneity in study design, participant age, OM diagnosis, and laboratory methods (Table 2). Where there were sufficient data to calculate I^2^; most were >70%, indicating moderate-high heterogeneity (Figure 2-4).

**Figure 2:**
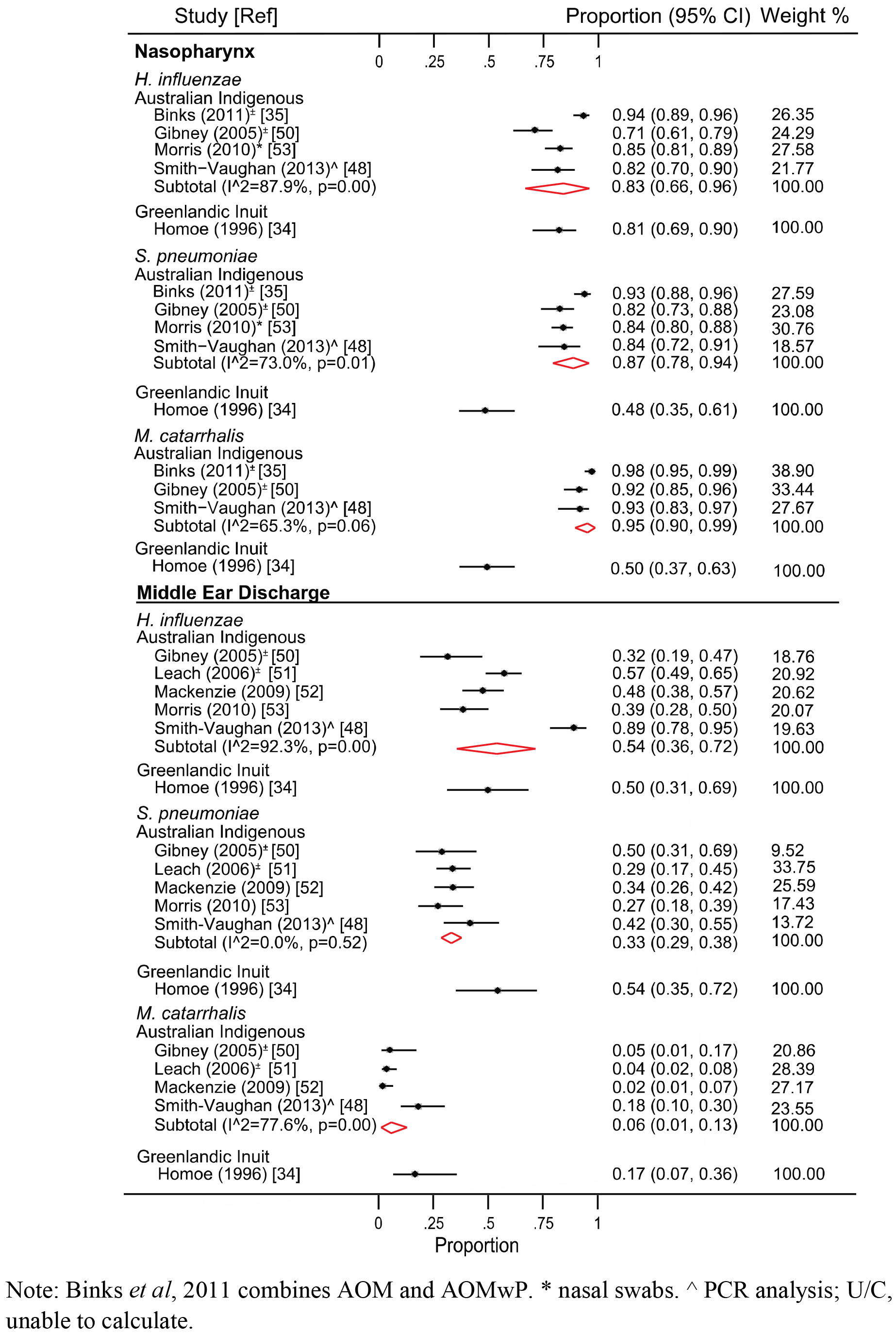
Bacteriology in relation to acute otitis media.

**Table 2:**
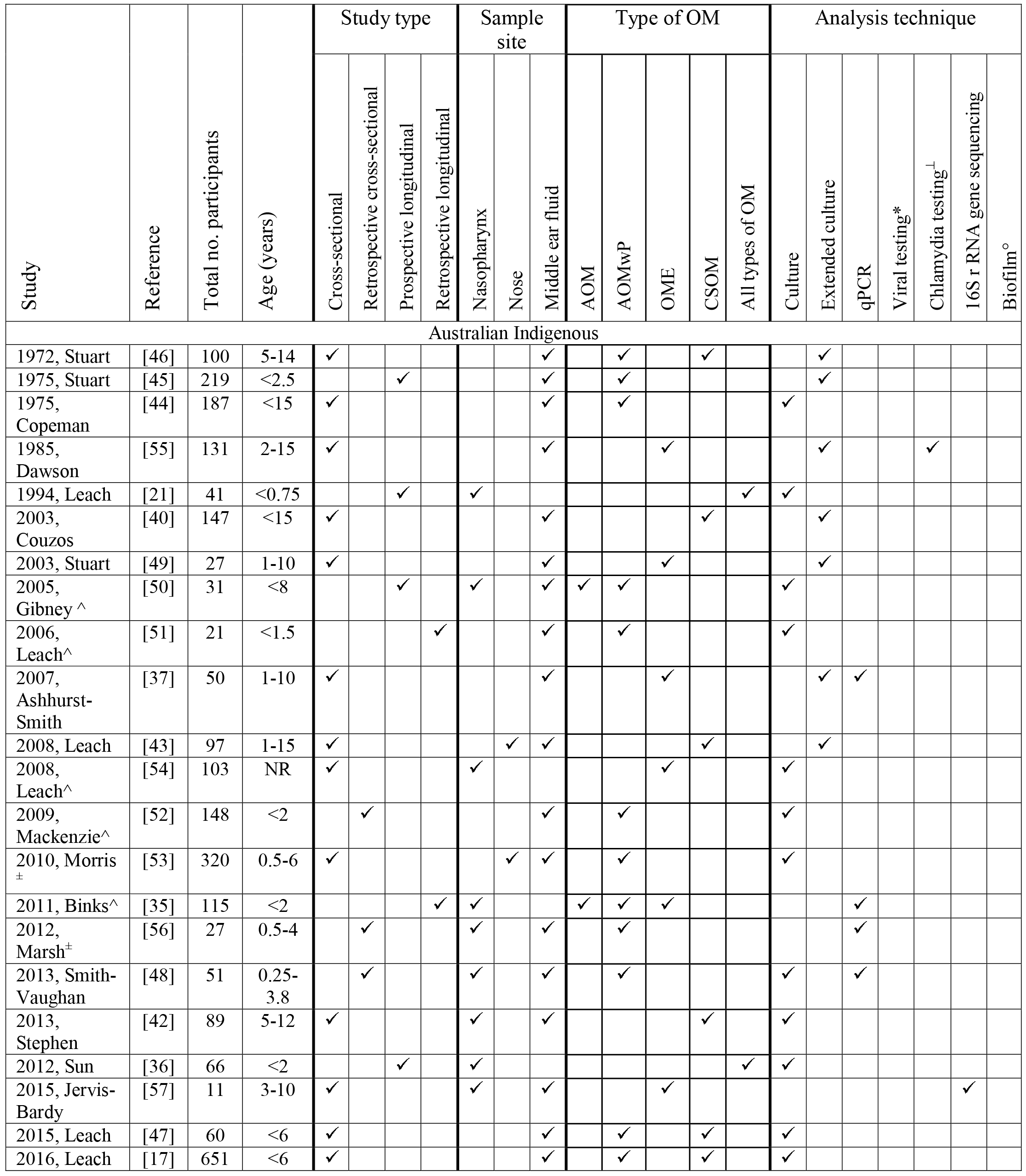
Characteristics of included studies.

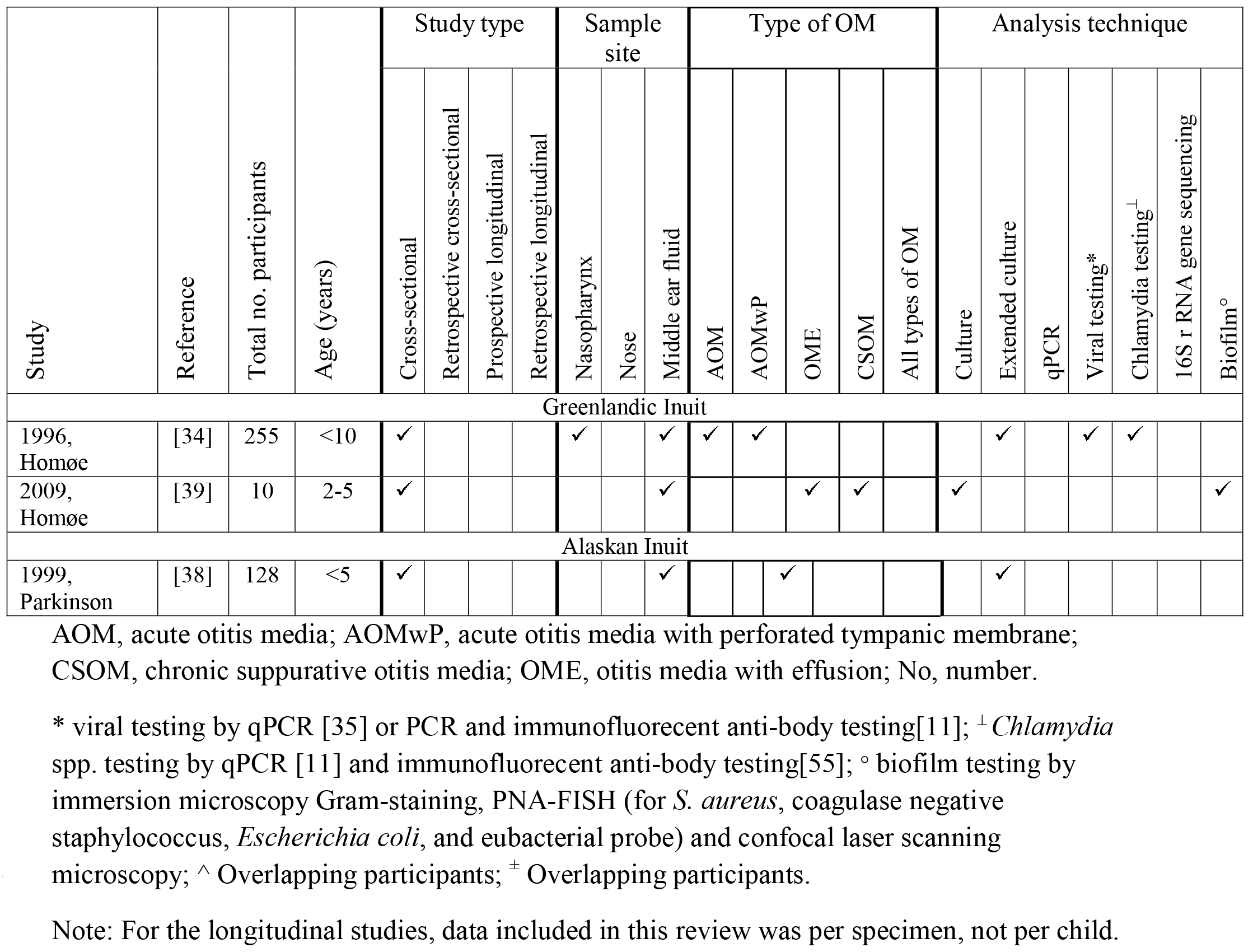

### OM clinical definitions and diagnosis

OM definitions used by the studies are outlined in Additional File 2. Acute OM (AOM) definitions were consistently based on otoscopy and tympanometry. OM with effusion (OME) was diagnosed based on a type-B tympanogram in 5/8 studies; the remaining three studies [37–39] reported data from intra-operative middle ear effusion (MEE) samples, without specifying OME diagnostic criteria. CSOM definitions were heterogeneous and included otorrhoea for >2 weeks [40, 41], >6 weeks [42], and broad descriptive terms [39, 43]. Three studies did not describe specific OM diagnostic criteria [44–46].

### Laboratory methods

Methods used to assess URT and middle ear bacteriology varied across studies (Table 2). Most studies (13/25) used culture conditions specific for detection of the main otopathogens. Nine studies used extended culture to detect a wider range of bacteria. For the culture-based studies, methodological details varied. Most culture-based studies (13/22) described the agar plates used and growth conditions [17, 21, 34, 36–40, 42, 43, 47–49]; however, reporting of phenotypic isolate identification tests varied. The remaining studies used non-specific terms or referred to other papers [44–46, 50–55].

Three studies used only molecular methods: two used species-specific qPCR targeting the main otopathogens or *Alloiococcus otitidis* [35, 56], and one used 16S rRNA gene sequencing [57]. One study used both culture and qPCR [48]. The three studies using qPCR [35, 48, 56] used the same gene targets for *S. pneumoniae* and *M. catarrhalis.* Two studies used the *hpd* gene to detect *H*. *influenzae* [35, 56] while another used an alternative gene target, *hpd3* [48]. Only one paper used qPCR to detect *A. otitidis* [56].

### Bacteriology

#### Acute otitis media

AOM bacteriology was reported for Australian and Greenlandic indigenous children, with high prevalence of the three main otopathogens in NP/nose and middle ear specimens across both populations (Figure 2 and Additional File 3). Co-infection with >1 otopathogen was common in the NP, although less frequent in MED (Additional File 3). NP colonization by *S. pneumoniae* (both populations) or *M. catarrhalis* (Australian Indigenous) was significantly related to AOM when compared to indigenous peers without OM [34, 35]. Beyond the main otopathogens, *A. otitidis, Staphylococcus* spp. and β hemolytic streptococcus were also detected in the middle ear discharge (MED) of children with AOM with perforated tympanic membrane (AOMwP) (Additional File 4).

#### Otitis media with effusion

The one study investigating NP microbiota, and all but one study exploring MEE in children with OME were from Australian Indigenous children. The three main otopathogens were highly prevalent in the NP in children with OME (Figure 3 and Additional File 3), although only *S. pneumoniae* and *M. catarrhalis* were significantly related to OME in the one study that included a control group [35]. Culture-based studies reported a low prevalence of otopathogens in MEE (Figure 3, Additional File 3); however, much higher rates were detected in the single study that used molecular methods [57] (Figure 3). Other bacteria detected in MEE by extended culture included *A. otitidis, Corynebacterium* spp., *Pseudomonas aeruginosa* and *S. aureus* (Additional File 4). The single 16S rRNA gene sequencing analysis (Australian Indigenous children) [57] found high rates of the genera *Dolosigranulum, Moraxella, Haemophilus* and *Streptococcus* (Mitis group) in the NP, and *Alloiococcus, Haemophilus* and *Corynebacterium* in MEE (Additional File 5)

**Figure 3:**
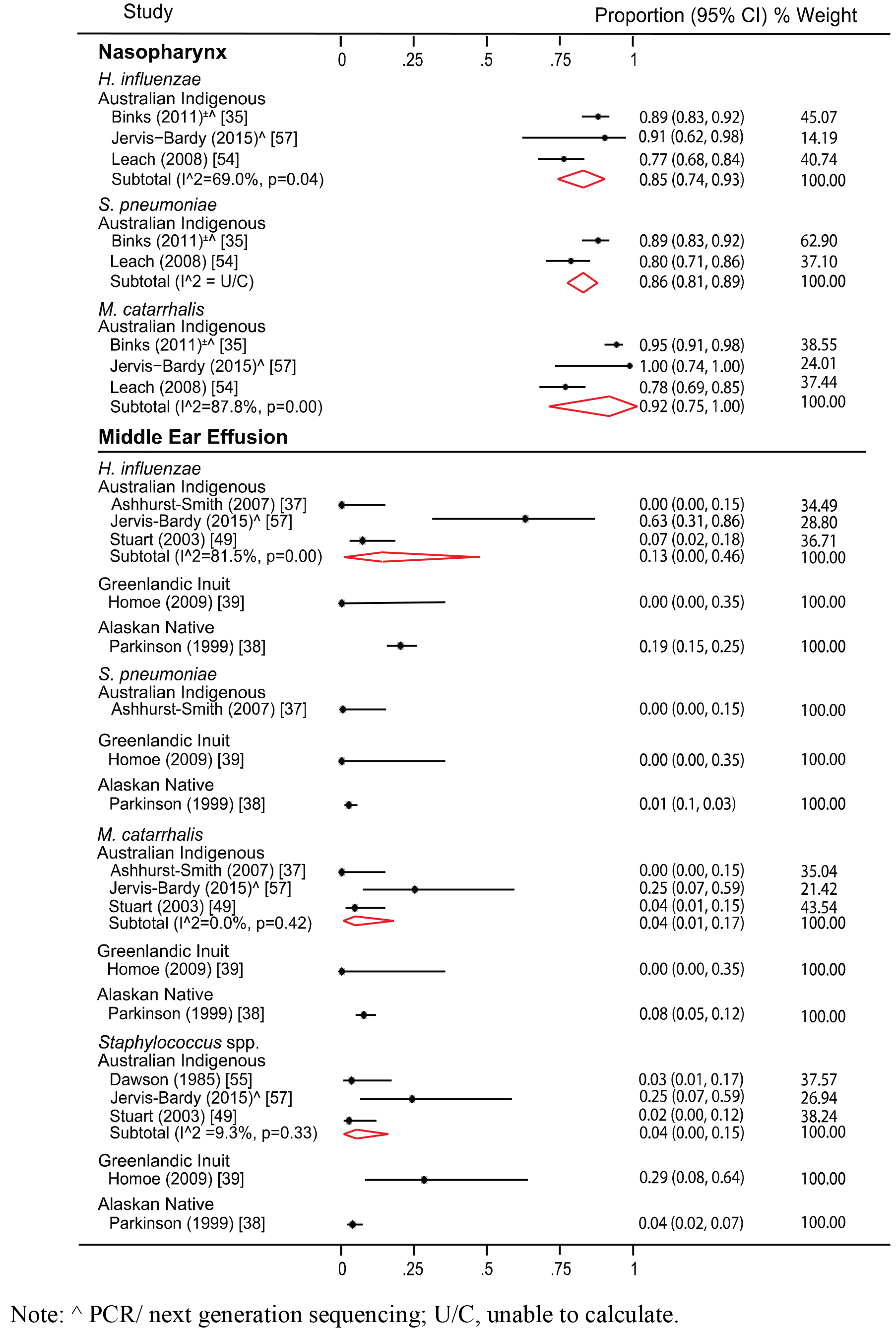
Bacteriology in relation to otitis media with effusion.

#### Chronic suppurative otitis media

All but one study investigating CSOM were from Australian Indigenous children. The most commonly reported bacteria from culture-based studies of MED from children with CSOM were *P. aeruginosa, S. aureus*, and *H. influenzae* (Figure 4). *P. aeruginosa* and *H. influenzae* were often detected in Australian Indigenous children, but not in the single study of Greenlandic Inuit children (Figure 4). Yeasts were reported in two Australian Indigenous studies (Additional File 4); one study [40] only detected *Candida*, *Aspergillus*, *Fusarium*, *Alternaria*, *Rhodotorula*, *Auerobasidium* or *Acrinomium* in 5% of MED samples. The other study [43] did not identify or specify the yeasts or fungi detected. No study used molecular methods to explore the URT or middle ear microbiota in CSOM.

**Figure 4:**
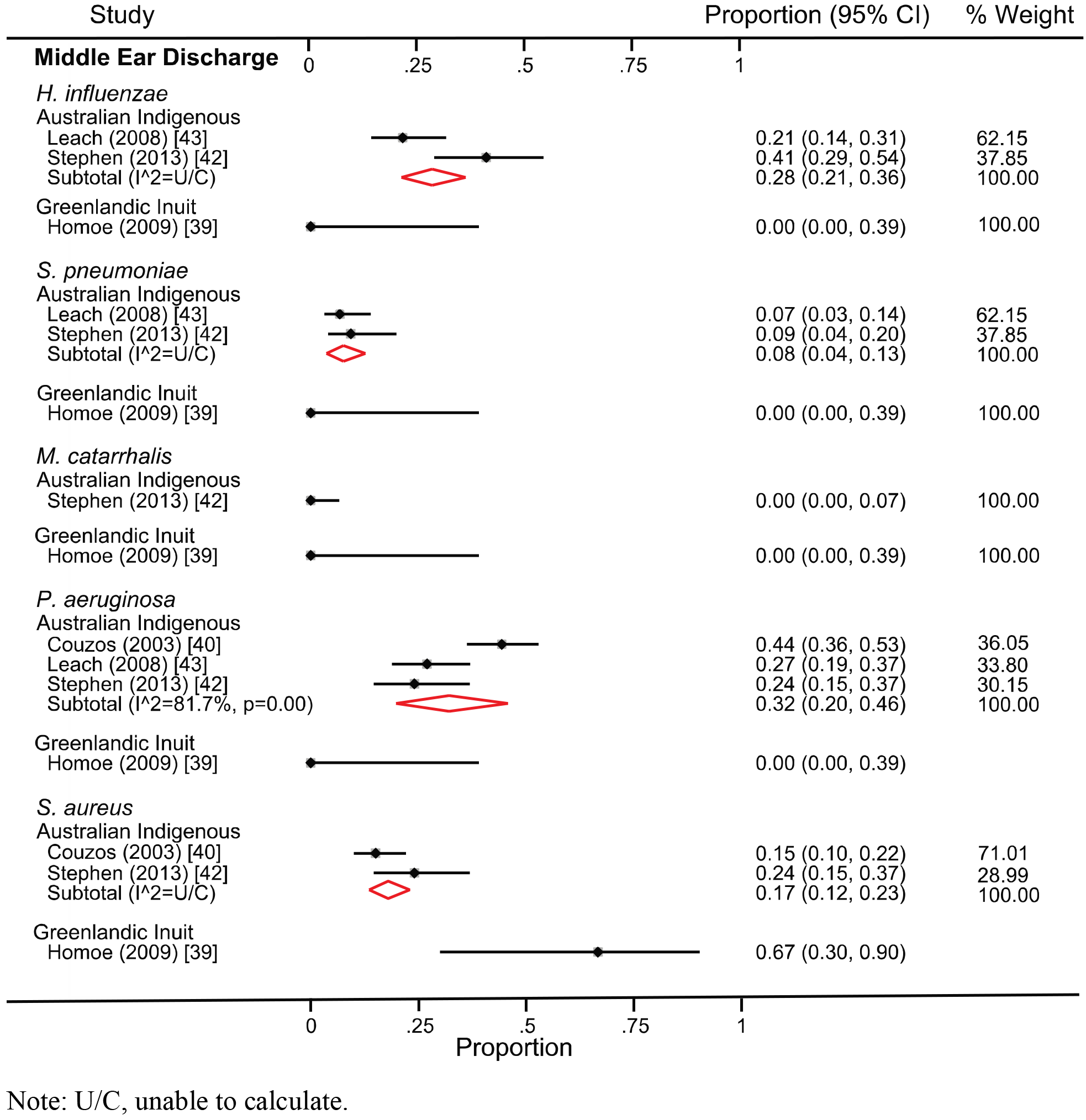
Bacteriology in relation to chronic suppurative otitis media.

**Figure 5:**
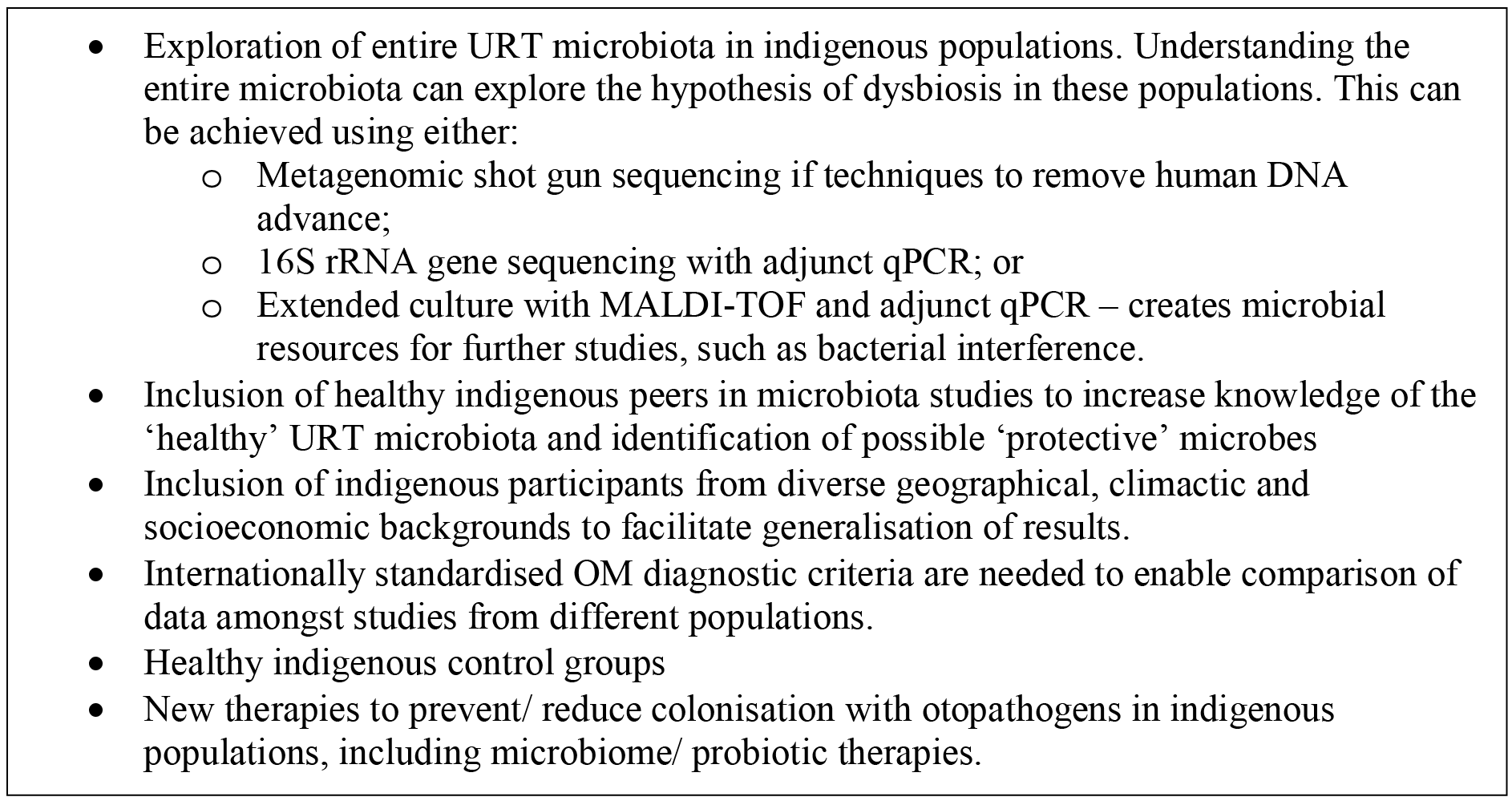
Recommendations for future research of OM microbiology in indigenous children.

### Nasopharyngeal carriage as a risk factor for otitis media

Two prospective cohort studies in Australian Indigenous children explored NP carriage of the three main otopathogens as a risk factor for OM (all types) [21, 36]. A birth cohort study by Leach *et al*. found that 31/36 (86%) children with their first episode of OM were colonized with at least one otopathogen [21]. This relationship between NP colonization and OM was stronger when >1 otopathogen was detected in the NP (odds ratio (OR) = 33.6, 95% CI 7.9 to 144) [21]. More recently, Sun *et al*. found that in Australian Indigenous children, early colonization (1 to <3 months of age) with *H. influenzae* was associated with OM in the first two years of life (OR = 3.71, 95% CI 1.22 to 11.23) [36]. All children (100%) who carried *H. influenzae* with either of the other main otopathogens were subsequently diagnosed with OM [36].

### Virology

One Australian [35] and one Greenlandic study [34] tested for viruses in children with OM (Additional File 4). These studies used different methods for viral detection and, aside from rhinovirus, tested for different viruses (Table 2). In Indigenous Australian children, only adenovirus in the NP was related to AOM (19%) and AOMwP (20%) compared to control children (6%) [35]. There was no relationship between the detection of viruses in the NP and OME [35]. In Greenlandic Inuit children, enteroviruses, rhinoviruses or ‘unspecified virus’ in the NP was related to AOM, compared to controls [34]. No studies tested for viruses in middle ear specimens in AOMwP, OME, or CSOM.

### Biofilm

One Greenlandic study used PNA-FISH to test for biofilm in middle ear specimens from children with CSOM or OME (obtained via sterile aspiration) [39]. Biofilm was detected in 5/6 (83%) MED samples from children with CSOM using a Eubacterial probe, but not in MEE from seven children with OME [39]. Further testing with species-specific probes found most biofilms (66%) contained *S. aureus.* One further sample contained a *Stenotrophomonas maltophilia* biofilm. For *S. aureus* and *S. maltophilia*, there was a 100% agreement between culture, Gram-staining and PNA-FISH results [39]. Species-specific probes targeting the main otopathogens were not tested.

## Discussion

This systematic review found the NP of most indigenous children with OM were colonized with the main otopathogens, particularly those with AOM. In contrast, children with CSOM demonstrate a different middle ear microbial profile compared to children with AOM and OME. Beyond the typical culturable bacteria, data are sparse, limiting our understanding of how the broader microbiota of the URT may contribute to OM pathogenesis and persistence in indigenous populations. The entire OM microbiome across all indigenous populations needs further investigation.

Our analysis highlights the important role of *S. pneumoniae* and *H. influenzae* in the pathogenesis of AOM/AOMwP and OME across indigenous populations, consistent with data from non-indigenous populations [58]. These otopathogens were detected at low rates in middle ear samples from children with AOMwP and OME; however, when molecular techniques were employed detection rates were much higher, particularly for *H. influenzae* [57], consistent with the increased sensitivity of molecular methods compared to culture [56, 59]. This suggests that current data, which are predominantly culture-based, may underestimate the prevalence of otopathogen colonisation in middle ear samples from indigenous children.

A different pathogen profile was reported from children with CSOM, including, *P. aeruginosa*, *S. aureus, H. influenzae* and fungi/ yeasts. Commensurate with this result, culture-based literature from non-indigenous children with CSOM often report *P. aeruginosa* and *S. aureus* in MED [60–65]. 16S rRNA gene sequencing of MED from children and adults with CSOM in New Zealand further detected *Alloiococcus* and *Streptococcus* [66]. In the CSOM studies included in this review, *Alloiococcus* would not have been detected, if present, as the specialist culture conditions or PCR required to detect this species were not used. The chronic perforation of the tympanic membrane in CSOM may allow for secondary infection of the middle ear by microbes present in the external auditory canal and could account for the different microbial profile compared to other types of OM. Confirming this; however, is difficult, particularly where a child has had prolonged otorrhoea with ear discharge draining into the canal. Sampling the canal flora of children with intact tympanic membranes as a comparison, may provide a solution.

Biofilms have been reported in middle ear specimens from non-indigenous children with CSOM and OME [41, 59, 67–69]; however, this systematic review uncovered very little data pertaining to biofilm in relation to OM in indigenous children. Considering the high rates of chronic OM, particularly CSOM, this is a noteworthy deficit of the literature.

This systematic review suggests other microbes, beyond the main otopathogens, may be contributing to OM in indigenous populations; however, there are few data relating to these taxa. Furthermore, detection of these microbes can require specific laboratory techniques. For example, *A. otitidis* detection requires extended culture methods [37] or molecular methods [56, 57]. Where these methods have been used, *A. otitidis* was commonly detected [37, 56, 57]; however, it remains controversial whether detection of this species is associated with the middle ear infection or specimen contamination by canal flora [56]. Viruses were seldom investigated in the included studies, and when investigated, different viruses were sought, and detection methods varied. Furthermore, only NP specimens were tested. Viruses are likely to play an important role in OM pathogenesis [70], through numerous potential mechanisms including altering the host immune response [71] and reducing response to antibiotic therapy [72]. Further research is required to determine the contribution of respiratory viruses in OM pathogenesis.

### Limitations of the current literature

The current literature is limited by methodological heterogeneity, in both the types of laboratory methods used and the OM definitions and diagnoses. The greatest source of methodological heterogeneity was the diversity of methods used to analyze the samples with varying specificities and sensitivities. Inconsistencies in OM definitions and diagnoses were most apparent in the CSOM data, reflecting the absence of internationally accepted definitions [73]. Other OM diagnoses were more consistent, largely because the data was published from a limited number of research groups. International guidelines on OM definitions, diagnosis and investigation of URT/ middle ear microbiota are needed. This will allow for more meaningful comparison of studies from around the world and facilitate future meta-analysis.

The quality of the data included in this review is impacted by the absence of healthy indigenous controls; limited information on participant recruitment; poor consideration of confounding variables; multiple studies where the microbiology is not the primary aim of the study; and population overlap. The absence of healthy indigenous control children may reflect the high burden of disease in many of these populations, for example <10% of Australian Indigenous children living in remote areas have healthy ears [7]. To establish a ‘OM microbiota’, comparison with healthy indigenous peers is required. Similarly, if samples from the external auditory canal are included when analyzing middle ear specimens, we may be able to delineate the role of microbes as contaminate, pathogen or secondary pathogen (e.g. *A. otitidis).* There was significant population overlap and small geographical area of recruitment for many studies in Australian Indigenous children. There is documented discordance in OM burden and prevalence of otopathogen colonization between urban and remote Australian Indigenous children [74, 75]. Therefore, this limited area of recruitment may impact on generalization of results across Australian Indigenous children.

### Future directions

To further our understanding of OM pathogenesis in indigenous populations, and to build upon the current pathogen-based disease model, further research is required to investigate the vast array of microbes that can occupy the URT, and how they relate to the known otopathogens to cause disease. The inclusion of healthy indigenous peers is vital to this goal. Identification of a ‘healthy’ microbiota in indigenous populations may uncover ‘protective’ microbes that can be developed into microbiome/ probiotic therapies to protect children from OM. To achieve this outcome, next generation sequencing can enable deeper exploration of the microbiota without *a priori* assumptions about the underlying bacterial community, which is required to guide culture-based methods. 16S rRNA gene sequencing, although limited by poor resolution at the species-level, can be augmented by qPCR to provide species-level identification [76]; however, this requires *a priori* assumptions about the bacteria that should be targeted. Likewise, qPCR for specific viruses is limited by *a priori* assumptions. These limitations may be overcome with metagenomic shotgun sequencing if the method can be optimized to overcome the technical limitations related to high proportions of human DNA in middle ear specimens. Alternatively, extended culture with MALDI-TOF can be used to provide a broader analysis of the microbiota to the species level [77] and has the benefit of providing material for further studies, such as bacterial interference studies.

## Conclusions

The URT microbiology in OM is highly complex and dynamic. Through this systematic review we demonstrated that the three main otopathogens are important in the pathogenesis of AOM across the indigenous populations included, and in non-indigenous peers. There is; however, a vast community of microbes present in the URT. How these microbes interact to promote or, perhaps more importantly protect, indigenous children from OM requires further investigation. With a more wholistic understanding of the microbial pathogenesis of OM in indigenous populations, we will be better equipped to develop new methods to prevent and treat OM in these populations.

## Competing interests

The authors declare that they have no competing interests.

## Funding

Coleman received support from an Avant Doctors in Training Research Scholarship, an NHMRC Postgraduate Research Scholarship (APP1133366) and a Queensland Health Junior Doctor Fellowship.

Marsh is supported by the NHMRC CRE in Respiratory Health of Aboriginal and Torres Strait Islander Children, grant number 1040830.

Cervin is support by the University of Queensland Faculty of Medicine Strategic Funding and The Garnett Passe & Rodney Williams Memorial Foundation.

